# Global assessment of *Mycobacterium avium* subspecies *hominissuis* genetic requirement for growth and virulence

**DOI:** 10.1101/568477

**Authors:** Marte S. Dragset, Thomas R. Ioerger, Maja Loevenich, Markus Haug, Niruja Sivakumar, Anne Marstad, Pere Joan Cardona, Geir Klinkenberg, Eric J. Rubin, Magnus Steigedal, Trude H. Flo

## Abstract

Nontuberculous mycobacterial infections caused by the opportunistic pathogen *Mycobacterium avium* subsp. *hominissuis* (MAH) are currently receiving renewed attention due to increased incidence combined with difficult treatment. Insights into the disease-causing mechanisms of this species have been hampered by difficulties in genetic manipulation of the bacteria. Here, we identified and sequenced a highly transformable, virulent MAH clinical isolate susceptible to high-density transposon mutagenesis, facilitating global gene disruption and subsequent investigation of MAH gene function. By transposon insertion sequencing (TnSeq) of this strain, we defined the MAH genome-wide genetic requirement for virulence and *in vitro* growth, and organized ~3500 identified transposon mutants for hypothesis-driven research. The majority (71 %) of the genes we identified as essential for MAH *in vitro* had a growth-essential mutual ortholog in the related and highly virulent *M. tuberculosis* (*Mtb*). However, passaging our library through a mouse model of infection revealed a substantial number (54% of total hits) of novel virulence genes. Strikingly, > 97 % of the MAH virulence genes had a mutual ortholog in *Mtb*. Two of the three virulence genes specific to MAH (i.e. no *Mtb* mutual orthologs) were PPE proteins, a family of proteins unique to mycobacteria and highly associated with virulence. Finally, we validated novel genes as required for successful MAH infection; one encoding a probable MFS transporter and another a hypothetical protein located in immediate vicinity of six other identified virulence genes. In summary, we provide new, fundamental insights into the underlying genetic requirement of MAH for growth and host infection.

**Author summary:** Pulmonary disease caused by nontuberculous mycobacteria is increasing worldwide. The majority of these infections are caused by the *M. avium* complex (MAC), whereof >90% arise from *Mycobacterium avium* subsp. *hominissuis* (MAH). Treatment of MAH infections is currently difficult, with a combination of antibiotics given for at least 12 months. To control MAH by improved therapy, prevention and diagnostics, we need to understand the underlying mechanisms of infection. While genetic manipulation of pathogens is crucial to study pathogenesis, *M. avium* (*Mav*) has been found notoriously hard to engineer. Here, we identify an MAH strain highly susceptible to high-density transposon mutagenesis and transformation, facilitating genetic engineering and analysis of gene function. We provide crucial insights into this strain’s global genetic requirements for growth and infection. Surprisingly, we find that the vast majority of genes required for MAH growth and virulence (96% and 97%, respectively) have mutual orthologs in the tuberculosis-causing pathogen *M. tuberculosis* (*Mtb*). However, we also find growth and virulence genes specific to MAC species. Finally, we validate novel mycobacterial virulence factors that might serve as future drug targets for MAH-specific treatment, or translate to broader treatment of related mycobacterial diseases.

## Introduction

*Mycobacterium avium* complex (MAC) is a group of genetically related and ubiquitously distributed opportunistic mycobacteria that can cause nontuberculous infections collectively called MAC disease [1]. *M. avium* (*Mav*), one of the MAC species, has been classified into subspecies *avium*, *paratuberculosis, silvaticum* and *hominissuis* based on molecular characterizations, prevalent hosts and diseases caused [2, 3]. The latter, *M. avium* subsp. *hominissuis* (MAH), can infect humans and lead to pulmonary and disseminated disease, particularly in immunocompromised individuals [4]. MAH infections are currently hard to treat, with a combination of antibiotics typically given for at least 12 months [5]. Similar to its relative *M. tuberculosis* (*Mtb*), the causative agent of tuberculosis, MAH proliferates within macrophages by hijacking normal phagosomal trafficking, overcoming the host’s elimination strategies [6–12]. Mechanisms of infection may therefore partly be conserved between the two species. MAH lacks the Type VII ESX-1 secretion system crucial for full *Mtb* virulence [13], suggesting they also differ in virulence strategies. While *Mtb* is an obligate human pathogen in nature, with limited survival outside the host, *Mav* is environmental and opportunistic, found in a variety of niches (e.g. soil, fresh water, showerheads) and a range of prevalent hosts [3]. MAH isolates exhibit high genetic variation [14], perhaps as an adaption to diverse niches and hosts. It is currently not known to what degree MAH and *Mtb* depend on similar mechanisms for growth and virulence, given the same selective conditions. Even so, MAH genes encoding factors required for basic proliferation and virulence may be attractive targets for improved MAH therapies, and may translate to the treatment of related mycobacterial diseases.

Transposon insertion sequencing (TnSeq), which combines transposon mutagenesis and massive parallel sequencing, has been widely used to determine the conditional requirement of bacterial genes on a genome-wide scale [15]. By massive parallel sequencing of libraries consisting of more than 100 000 transposon mutants, the genetic requirement of *Mtb*, *M. marinum* and *M. avium* subsp. *paratuberculosis* has been defined, for *in vitro* growth or during infection [16–20]. For MAH, transposon mutagenesis has also been of great importance to identify virulence genes [21–28]. However, studies in MAH have to date been limited in library sizes, identifying a smaller number of potential virulence genes per screen. The majority of past MAH mutant libraries were constructed using the clinical isolate MAH 104 [21–25, 28]. 104 was the first MAH strain with a publicly available, fully assembled genome [29], and has thus naturally been widely studied, also by us (exemplified in [8, 10, 11, 29, 30]). MAH 104 is, however, resistant to transposon mutagenesis by ϕMycoMarT7, the phagemid preferred to generate high-density mycobacterial libraries [16–20, 31, 32]. Moreover, MAH 104 (along with many other *Mav* strains) transforms with low efficiency [28, 33–35], complicating the use of this strain in hypothesis-driven genetic manipulation. To facilitate genome-wide investigation of MAH gene function, we identified an MAH strain (MAH 11) highly susceptible to both ϕMycoMarT7-mediated transposon mutagenesis and transformation. We used this strain to generate a transposon insertion library with ~66% saturation density, which we profiled for genes required *in vitro* and in a mouse model of infection. In fact, MAH 11 is currently used as a screening strain in mycobacterial drug discovery programs [36, 37], adding further value to determining the growth requirements of this particular strain. Finally, we constructed an ordered subset of our transposon library in 384-well format (with a unique clone in each well) and used a multiplex sequencing strategy to identify the disrupted gene in each well, providing access to ~3500 MAH mutants. With these, we validated novel MAH virulence factors. Fig 1 summarizes the overall experimental setup of our study.

**Fig 1:**
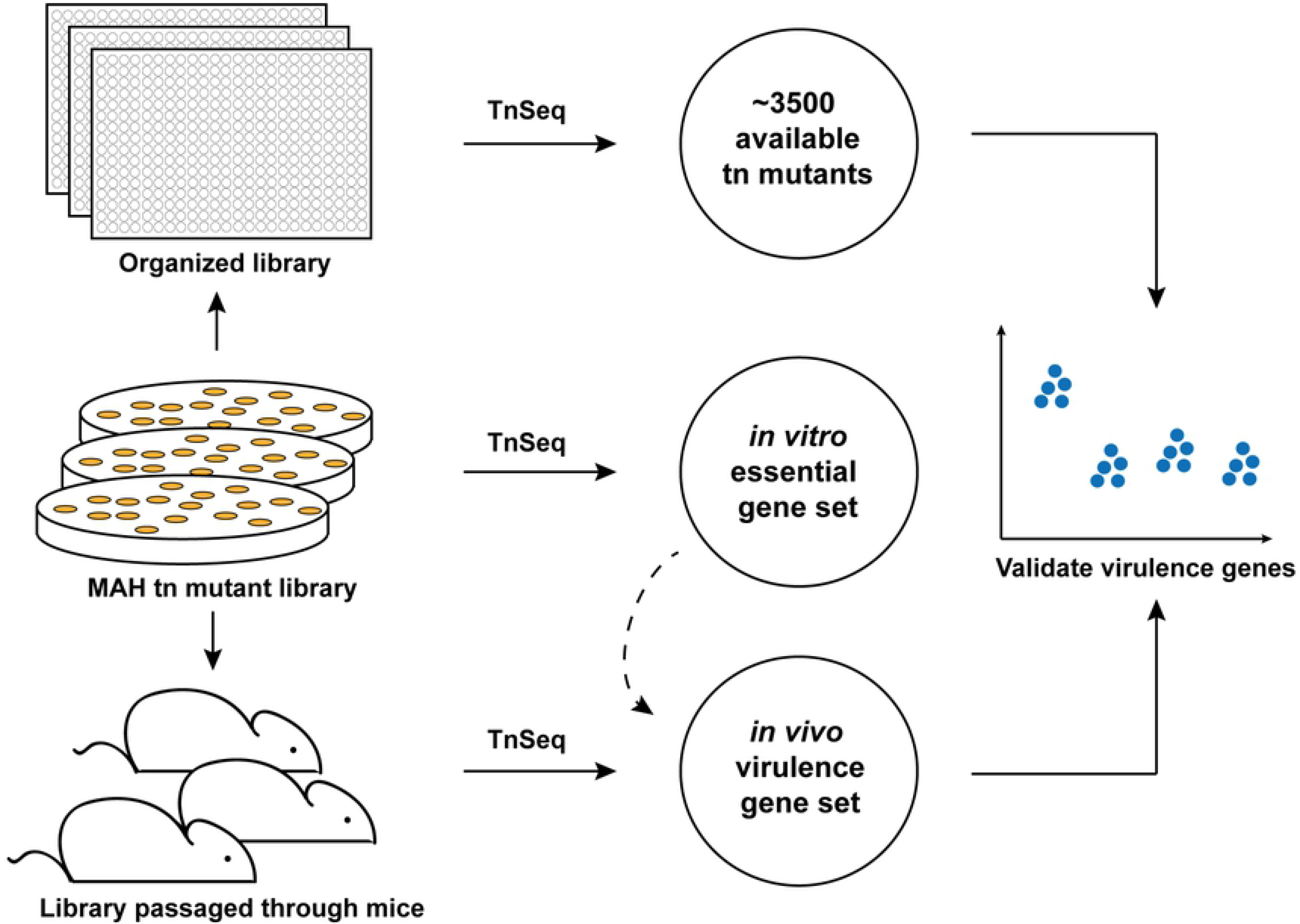
The experimental setup of the study. An MAH transposon mutant library of 170 000 mutants (~66% coverage) was generated by selection on 7H10 agar medium. This library was further subjected to mouse infection or organized mutant-by-mutant in 384-well plates (24 plates). By TnSeq of the *in vitro*-selected (7H10), *in vivo*-selected (C57BL/6 mice), and the organized library, the MAH essential- and virulence gene sets were defined, and the location of ~3500 organized mutants were identified. The output from the *in vitro* selected TnSeq’ed library was used to identify virulence genes (dashed line). Finally, a subset of virulence gene hits was validated by mouse infection experiments. Tn, transposon.

## Results

### Identification of a ϕMycoMarT7-transducible MAH strain

We aimed to find an MAH strain in which we could create high-density transposon mutant libraries. The transposon donor phagemid ϕMycoMarT7 is widely used (and recommended over Tn*5367* transposition [31]), for efficient mycobacterial transposon mutagenesis [16–20, 32]. ϕMycoMarT7 is derived from ϕAE87, which originates from mycobacteriophage TM4 [38, 39]. TM4 has been inconsistent in its ability to transduce various *Mav* strains and unable to infect the commonly studied genome sequenced strain MAH 104 [40]. In agreement with these observations, we failed to obtain kanamycin resistant mutants (marker for successful transposition of the ϕMycoMarT7-encoded *Himar1* transposon) when attempting to transduce MAH 104. Hence, to identify a ϕMycoMarT7-transducible strain of MAH, we screened seven in-house clinical *Mav* isolates originating from patients at the National Taiwan University Hospital, Taiwan. Around 70% of the strains resulted in kanamycin resistant colonies after transduction, indicating *Himar1* transposition. Based on particularly efficient transducibility (up to 300 000 kanamycin-resistant colonies per ml starting culture in our initial small-scale screen) and general ease to handle, we focused on a strain isolated from an HIV positive patient’s bone marrow, MAH 11. By PCR, we confirmed the presence of transposon inserted in 10 out of 10 colonies tested (S1 Fig). Hence, this MAH strain is highly susceptible to ϕMycoMarT7-mediated transposon mutagenesis.

### MAH 11 genome sequence

We sequenced the genome of the ϕMycoMarT7-transducible MAH 11 strain on an Illumina HiSeq 2500 instrument in paired-end mode with a read length of 125 bp, yielding a mean depth of coverage of 55.4. The sequence was assembled by a comparative assembly strategy, using MAH 104 as a reference sequence, augmented with contig-building to build large-scale indels. The Illumina data was supplemented with long reads (up to 40kb) from a PacBio sequencer, which were used to confirm the connectivity of the chromosome. The length of the genome is 5,105,085 bp, and is GC-rich (69.2%) like that of other mycobacteria (Fig 2A). The genome sequence is highly concordant with the recently reported draft genome of MAH 11 (70 contigs) [41]. The primary discrepancies involve repetitive sequence elements, such as differences in locations of native transposons and copy-numbers of MIRU tandem repeats, for which assembly from short reads can be ambiguous.

**Fig 2:**
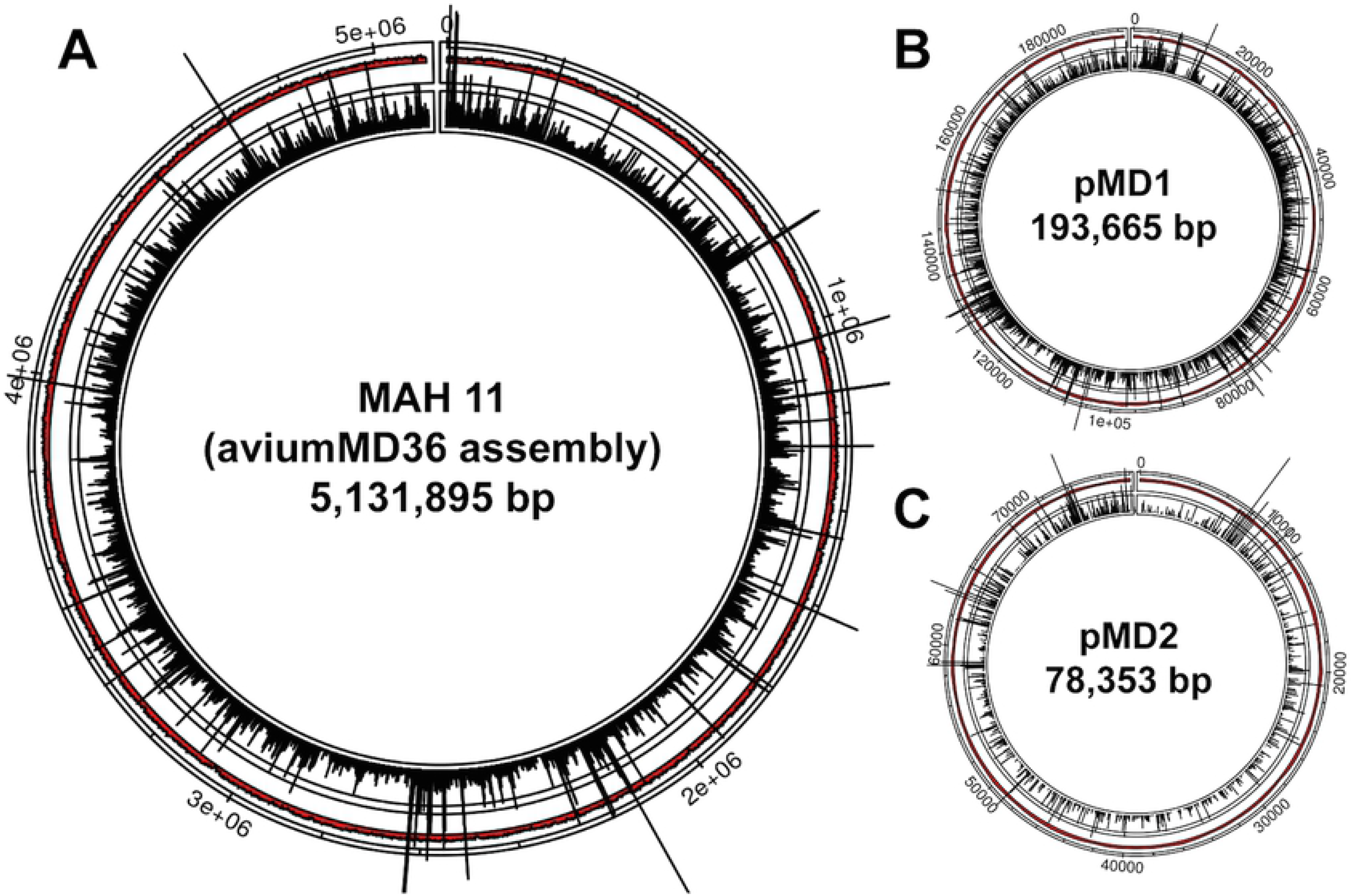
Distribution of transposon insertion counts. Transposon insertion counts across the MAH 11 genome (A) and the two plasmids, pMD1 (B) and pMD2 (C). The height of the black bars represents the number of insertion counts at the respective genome site.

MAH 11 is positively identified as *M. avium* subsp. *hominissuis*, based on 16S rRNA and *hsp65* sequences that are identical to MAH 104 (but distinct from other subspecies, like *M. avium* subsp*. avium*) [2]. However, relative to MAH 104, there are substantial numbers of SNPs (approximately 10 SNPs per 1kb) and a cumulative loss of ~377 kb, showing it is a distinct lineage (Fig 3A). The reductions are clustered in several large-scale deletions, the largest of which (>40kb) are listed in Fig 3B. This variability has been seen in other *Mav* isolates [42, 43], and several of the large-scale deletions correspond to known large-scale polymorphisms [42]. In addition, there are several large-scale insertions (Fig. 3B), including a prophage (56 genes, *b6k05_17725*-*18015*, inserted at coordinate 3.77 Mbp). This phage is almost identical to a prophage previously reported in the genome of *M. chimaera* str. MC045 (RefSeq NZ_LT703505.1). Also, a cluster of 48 non-phage metabolic genes (*b6k05_03885*-*04170*; 57kb) is inserted in the tail of existing MAH 104 prophage phiMAV-1 (*mav_0779-0841*; [44]), containing a variety of hydrolases, reductases, dehydrogenases, and monooxygenases. This gene cluster has been previously observed in other MAH strains, e.g. strain H87 [45]. The MAH 11 genome contains at least 14 copies of IS1245 (similar to MAH 104; [46]), but none of IS901 (associated with members of the MAC complex primarily infecting birds; [42, 47]). S2 Fig shows the position of MAH 11 in a phylogenetic tree (created using PHYLIP, http://evolution.genetics.washington.edu/phylip.html) together with 21 other *M. avium* genomes obtained from NCBI GenBank, including *M. avium* subsp. *paratuberculosis* (MAP) K10 and three *M. avium* subsp. *avium* (MAA) as outgroup strains.

**Fig 3:**
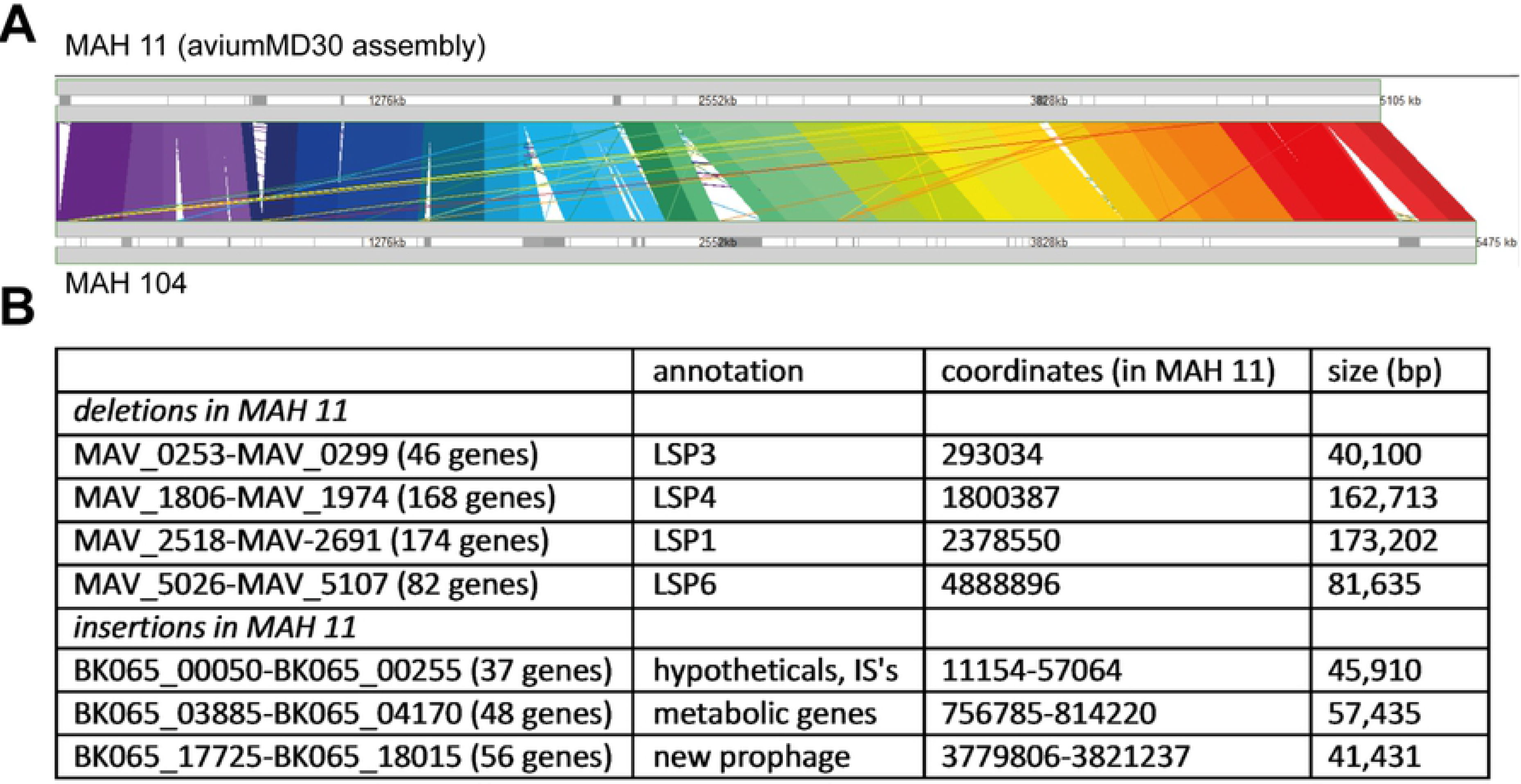
Synteny plot. (A) Plot of synteny between MAH MD (top) and MAV 104 (bottom), made with M-GCAT [81]. (B) The largest (>40kb) insertions and deletions in MAH 11 relative to MAH 104.

We identified 4653 ORFs (along with 1 copy of the rRNAs (16S, 23S, 5S) and 42 tRNAs, similar to MAH 104) using the NCBI Prokaryotic Genome Annotation Pipeline [48]. 4211 genes have mutual orthologs with MAH 104 (where each gene in one organism is the best match for the ortholog in the other organism, with BLAST E-value < 10^−10^, S1A Dataset). Almost all of these orthologs (4123) have ≥ 96% amino acid identity, and nearly half (1730) have 100% amino acid identity. For simplicity, we will, where applicable, refer to MAH ORFs using the MAH 104 locus tags (*mav_xxxx*) from here on. Compared to *Mtb* (H37Rv), MAH 11 has ~500 more genes, and over half the genes in each genome (2681) has a mutual ortholog in the other genome (S1B Dataset). Most of the remaining genes also have orthologs, but their E-values are above the stringent threshold of 10^−10^ (i.e. 1052 MAH genes have no clear *Mtb* ortholog, S1C Dataset), or their specific partner in the other genome is ambiguous, as is sometimes the case in duplicated gene families.

### MAH 11 plasmids

Two large extra-chromosomal contigs that appear to represent circular plasmids were detected. One, pMD1 (193kb, 162 ORFs) (Fig 2B and S1D Dataset), bears weak similarity (based on BLAST search) to parts of plasmids in a wide range of other mycobacteria. The other, pMD2 (78kb, 66 ORFs) (Fig 2C and S1E Dataset), is nearly identical to conjugative plasmid pMA100 from MAH strain 88Br (though reduced, since pMA100 is 116kb, S1F Dataset) [49], and bears partial similarity to the conjugative pRAW-like plasmids found in several slow-growing mycobacterial species [50].

### MAH 11 is highly transformable

*Mav* is notoriously hard to transform [28, 33–35], complicating introduction of new DNA and thus genetic engineering of this species. In our and others’ experience, MAH 104 is transformable with low efficiency [34, 51]. We investigated the transformation frequency of MAH 11 and found that this strain is around 100 times more susceptible to obtain plasmid DNA compared to the 104 strain, using optimized protocols for *Mav* electroporation (Fig 4A) [33]. We did not observe a notable difference in cell wall integrity (potentially explaining different susceptibility to electroporation) between MAH 11 and MAH 104 when subjecting them to sodium dodecyl sulfate (SDS) stress (S3 Fig). In summary, MAH 11 might be particularly apt for hypothesis-driven genetic approaches.

**Fig 4:**
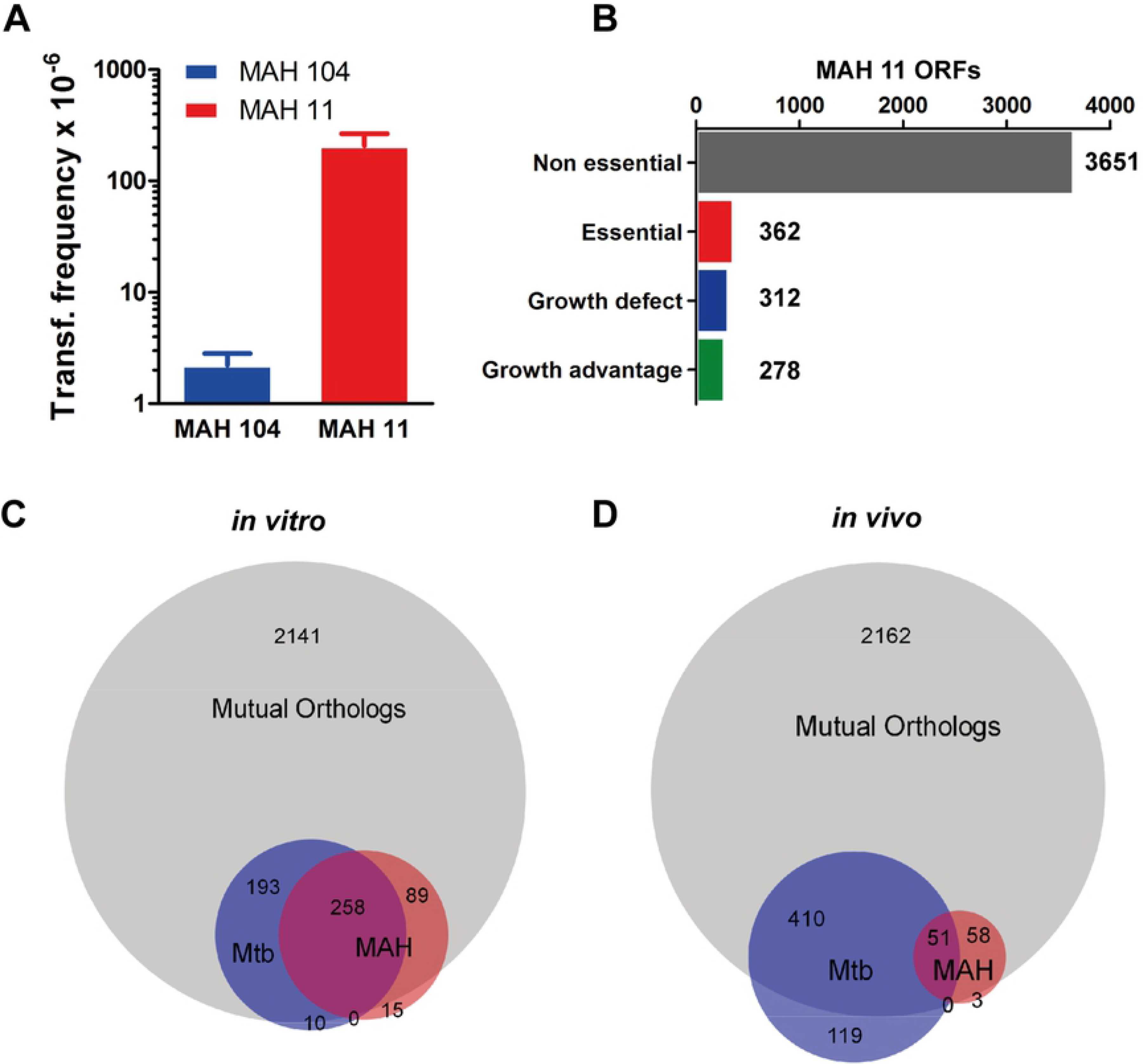
The MAH *in vitro* essential and *in vivo* virulence gene sets. (A) MAH 11 and MAH 104 transformation frequencies. Data show mean ± SEM of three individually electroporated samples. The result is representative of two independent experiments. (B) Number of MAH 11 genes defined as essential, growth defect, growth advantage and non-essential. (C) Venn diagram showing degree of overlap between MAH 11 and *Mtb* mutual orthologs (grey), MAH 11 *in vitro* essential genes as defined in this study (red), and *Mtb in vitro* essential genes (blue) as defined by DeJesus *et al*. [55]). (D) Venn diagram showing degree of overlap between MAH 11 and *Mtb* mutual orthologs (grey), MAH 11 virulence genes as defined in this study (red), and *Mtb* virulence genes (blue) as defined by Zhang *et al*. [16]. The Venn diagrams were created using Biovenn [82].

### MAH *in vitro* essential gene set

To define genes required for MAH *in vitro* growth we generated, by ϕMycoMarT7-mediated transduction, a library of ~170,000 transposon mutants selected on 7H10 medium. The *Himar1*-based mariner transposon of ϕMycoMarT7 inserts randomly at TA dinucleotides [52], of which there are 55,516 sites in the MAH 11 genome (excluding plasmids). We sequenced the transposon junctions of two independent libraries, mapped the genomic position of the transposon insertion sites (insertion counts), and counted insertions (reduced to unique templates using barcodes [53]) (S1 Table). The library had a saturation of 66.3%, with insertions at 36,813 out of 55,516 TA sites. By gene requirement analysis, using a Hidden Markov Model incorporated into the Transit platform [54], we defined 362 genes as essential for *in vitro* growth, 312 as genes causing growth-defect when disrupted, 278 as genes causing growth-advantage when disrupted and 3651 genes as non-essential for growth (Fig 4B, S1G Dataset). 71% (258/362) of MAH 11’s essential genes had an essential ortholog in *Mtb* (as defined by DeJesus *et al*. [55]). Remarkably, very few MAH (15) and *Mtb* (10) essential genes (~4% and ~2% of total essentials, respectively) did not have a mutual ortholog in the other species (Fig 4C, S1G Dataset), suggesting that the vast majority of genes required for *in vitro* proliferation are conserved between MAH and *Mtb*.

Almost all genes found on the two MAH 11 plasmids were non-essential for *in vitro* growth (S1H and S1I Datasets). However, on pMD1, four genes caused growth defects of MAH 11 when disrupted; two encoding hypothetical proteins, one homologous to a gene encoding chromosome partitioning protein, *parB*, and one homologous to DNA processing protein-encoding *dprA*. On pMD2, a gene homologous to *rep*, involved in plasmid replication, caused growth defect when disrupted. Taken together, we identified 674 chromosomal and five plasmid MAH genes that were essential or caused a growth defect *in vitro* when disrupted.

### MAH 11 establishes infection in mice

We and others have shown that MAH 104 is virulent in mice [8, 11, 12, 22]. We thus examined whether MAH 11 would be suitable to study the role of MAH genes *in vivo*. C57BL/6 mice were infected *intraperitoneally* with MAH 11 or MAH 104, and organ bacterial load was analyzed in the chronic phase of infection. As we have previously shown, bacterial loads remained relatively constant in liver and spleen from 22 to 50 days after initial MAH 104 infection [12]. The same trend was seen for MAH 11, albeit with an overall lower bacterial load compared to MAH 104, especially in the spleen (Fig 5A and B). MAH 11 and 104 grew comparably in 7H9 medium (Fig 5C), suggesting MAH 11 is less virulent than MAH 104 in mice.

**Fig 5:**
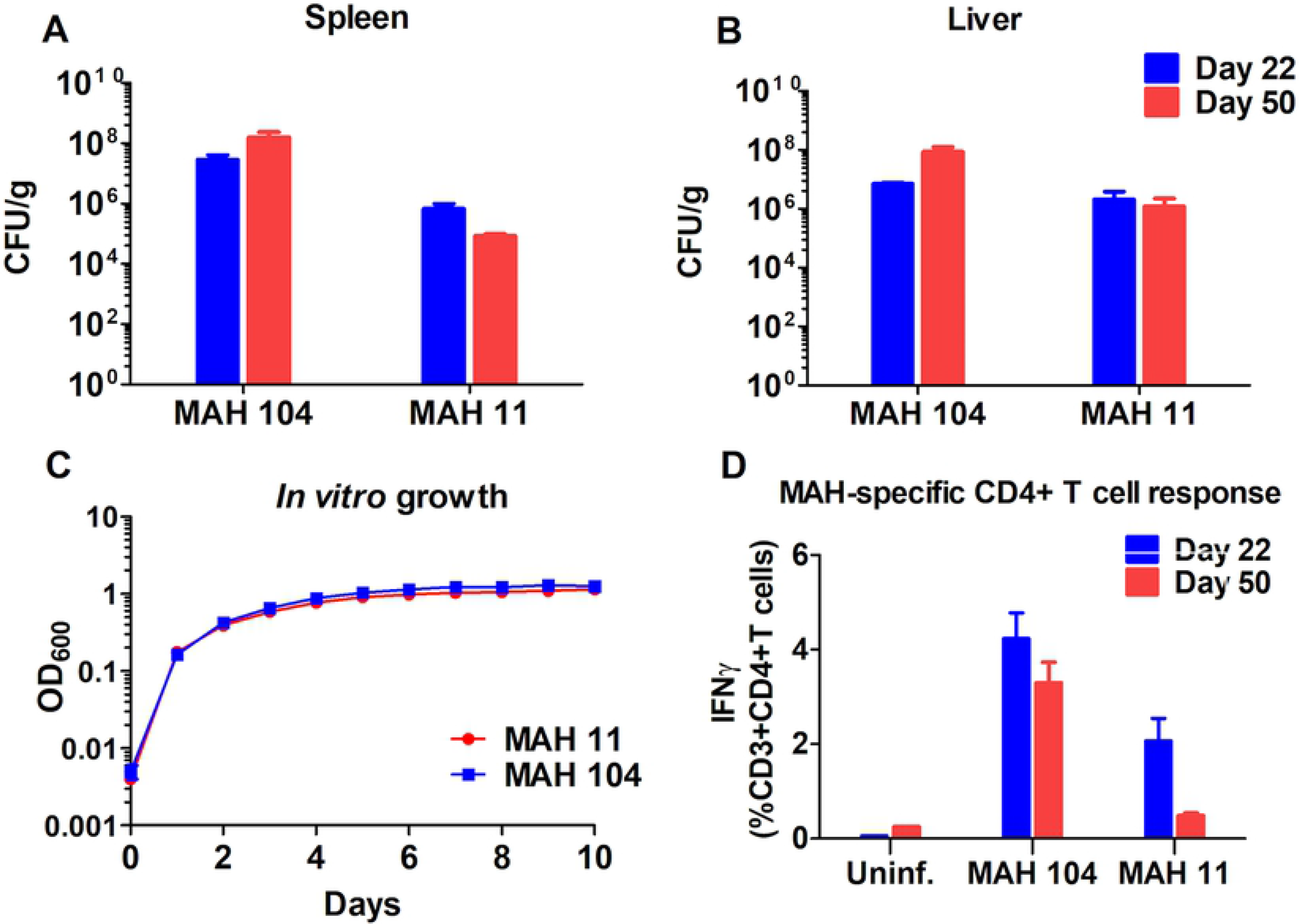
MAH 11 and MAH 104 mouse model infection. Bacterial burden in spleen (A) and liver (B) of mice after 22 and 50 days of infection with MAH 104 and MAH 11. Data show mean ± SEM of four infected mice in each group. (C) MAH 104 and MAH 11 *in vitro* growth (7H9 medium). Data show mean ± SEM of three replicate samples per condition. The result is representative of two independent experiments. (D) Frequencies of MAH-specific CD4+ T cells after 22 and 50 days of MAH 104 and MAH 11 infection. Splenocytes from all mice were stimulated overnight with MAH and frequencies of IFNγ-producing CD4+ effector T cells were analyzed by flow-cytometry. Data show mean ± SEM of four infected mice per group. Mice experiments are representative of two independent experiments (for 50 day time point).

T cells produce effector cytokines upon activation to elicit an adaptive immune response towards infections. To control *Mav* infection, production of IFNγ effector cytokine by CD4+ T helper 1 cells is of particular importance [56]. We have previously monitored anti-mycobacterial T cell responses to MAH 104 [12]. To investigate whether MAH 11 is suitable to study MAH-specific host immune responses, we measured mycobacteria-specific CD4+ T cell responses after mouse infection. Fig 5D shows frequencies of MAH-specific CD4+ T helper 1 cells producing IFNγ effector cytokine after infection. Interestingly, the frequencies of IFNγ-producing CD4+ T cells were found lower in MAH 11-infected mice compared to MAH 104-infected mice after 50 days of infection, possibly reflecting the lower organ bacterial loads (Fig 5A and B). Nevertheless, MAH 11 appears suitable to study mycobacterial disease mechanisms, as well as host responses, in a mouse infection model.

### MAH virulence gene set

To identify genes required for MAH virulence we infected six mice with our MAH library and analyzed bacterial load from the spleen and liver (organs from two animals each were pooled, resulting in three spleen libraries and three liver libraries) after 26 days of infection. We sequenced the harvested libraries, which yielded saturation of 61.9% and 71.0% (combined over replicates) for spleen and liver, respectively (S1 Table). We then defined the genetic requirement for infection using a Transit-incorporated resampling algorithm for comparative analysis [54], comparing output data from sequenced libraries before and after infection. We identified 144 and 128 genes as required for spleen and liver infection, respectively (S1B Dataset). 112 genes were required for survival in both organs (~80% overlap). Direct comparison of the spleen and liver datasets by resampling did not reveal any statistically significant differences, and hence no genes uniquely required for colonization of either organ were identified. Among the core genes identified (found in both spleen and liver) were 51 previously identified in *Mtb* mouse model TnSeq experiments [16], including well-established mycobacterial virulence genes like *uvrABC* (the UvrABC endonuclease complex [57]), *secA2* (alternative ATPase of Sec secretion pathway [58]), *icl* (isocitrate lyase [59]), *bioA* (within the biotin biosynthesis pathway [60]) and *glcb* (malate synthase [61]). However, importantly, we identified 61 core genes (92 genes with spleen and liver genes combined) not previously detected by *Mtb* TnSeq virulence gene screening (Fig. 4D)[16]. Some of these genes were found in genetic clusters, like six genes within the region encoding the Type VII ESX-5 secretion system. Other genes were found in operons (for instance *prcA*/*prcB*, *mav_3300*/*ripA*, *mav_3691*/*rbfA*) or in close genomic vicinity (*mav_4154*/*4158*/*4159*/*4160/4163*). Strikingly, only three (< 3%) core MAH virulence genes did not have a mutual ortholog in *Mtb* (Fig 4D). Two of them, *mav_4273* and *mav_4274*, are potentially co-expressed, and both encode proline-proline-glutamic acid (PPE) family proteins. PPE proteins might show ambiguity in ortholog matching due to large duplications within the family. However, by BLAST search, the two PPE proteins have clear orthologs in MAC species [62], but not in other well-known mycobacterial species like *Mtb*, *M. bovis*, *M. abscessus* nor *M. leprae* (albeit both PPE proteins show a weak similarity to *M. marinum* PPE14 (*mmar_1235*) with 53 and 57% amino acid identity, respectively). The last of the three MAH virulence genes without *Mtb* mutual orthologs, *mav_4409*, encodes a putative acyltransferase.

As could be expected, most genes found on the two MAH 11 plasmids were non-essential for infection (S1J-M Datasets); the only exceptions were an AAA-family ATPase on pMD1 (out of 162 ORFs), and two ORFs of unknown function on pMD2 (66 ORFs).

### Identification of mutants in an organized MAH library

Organized libraries of insertion mutants are of great value for hypothesis-driven research. Vandewalle *et al.* developed a method where sequence tagging transposon library pools was used to bulk-identify, by TnSeq, both the gene disrupted and the location of the mutant within an organized (plated) *M. bovis* BCG transposon library [63]. We employed such a strategy to identify mutants within an ordered MAH 11 library of 9216 transposon-insertion mutants, plated in 384-well plates using a colony-picking robot. We tagged the library by plates, rows and columns, then pooled and sequenced the samples, and analyzed the barcode-to-genome coordinate maps to determine the location of the mutants and the transposon insertion sites in bulk. We were able to map the specific location of 2696 unique transposon insertion mutants within 1697 (34.8%) of the 4881 MAH 11 ORFs (plasmid ORFs included) (S1N Dataset). Transposon insertion sites that mapped ambiguously to more than one plate, column and/or row were disregarded; these might be due to a relatively high number of mutant duplicates in the picked library. 3161 wells had a unique clone, and only 155 wells had more than one mutant assigned to them (S1O Dataset). The latter might be due to mutants clumping in colonies picked, incomplete sterilization of the robotic picking device between rounds of colony picking, or transposon insertions in repetitive or duplicated regions. To experimentally verify the correct location of mutants, we sequenced 11 mutants picked from 11 wells (S2 Table). All mutants had the transposon inserted at the location predicted by our TnSeq approach. Taken together, we identified the transposon insertion site and mapped the unambiguous location of 3489 clones (including those in intergenic regions), providing access to a plethora of mutants to study the role of the respective MAH genes.

### *uvrB* is required for MAH virulence

UvrABC is an enzyme complex involved in *Escherichia coli* nucleotide excision repair [64]. The genes encoding the mycobacterial homologues of the three members of the complex, *uvrA*, *uvrB*, and *uvrC*, were all defined as virulence genes in our screen. *uvrB* has previously been implicated in *Mtb* virulence, via protection against host-mediated reactive nitrogen and oxygen intermediates [57]. To verify the involvement of *uvrB* in MAH virulence, we infected mice with an available *uvrB* transposon mutant (*uvrB::tn*) and complemented mutant for 26 days. *uvrB::tn* showed reduced bacterial burden in infected mice compared to MAH 11 wild-type (wt) and complement-infected animals (Fig 6A and B). Neither the *uvrB* mutant nor the complemented mutant showed reduced fitness *in vitro* (Fig 6E), hence, our results suggest that UvrB is required for full virulence in MAH.

**Fig 6:**
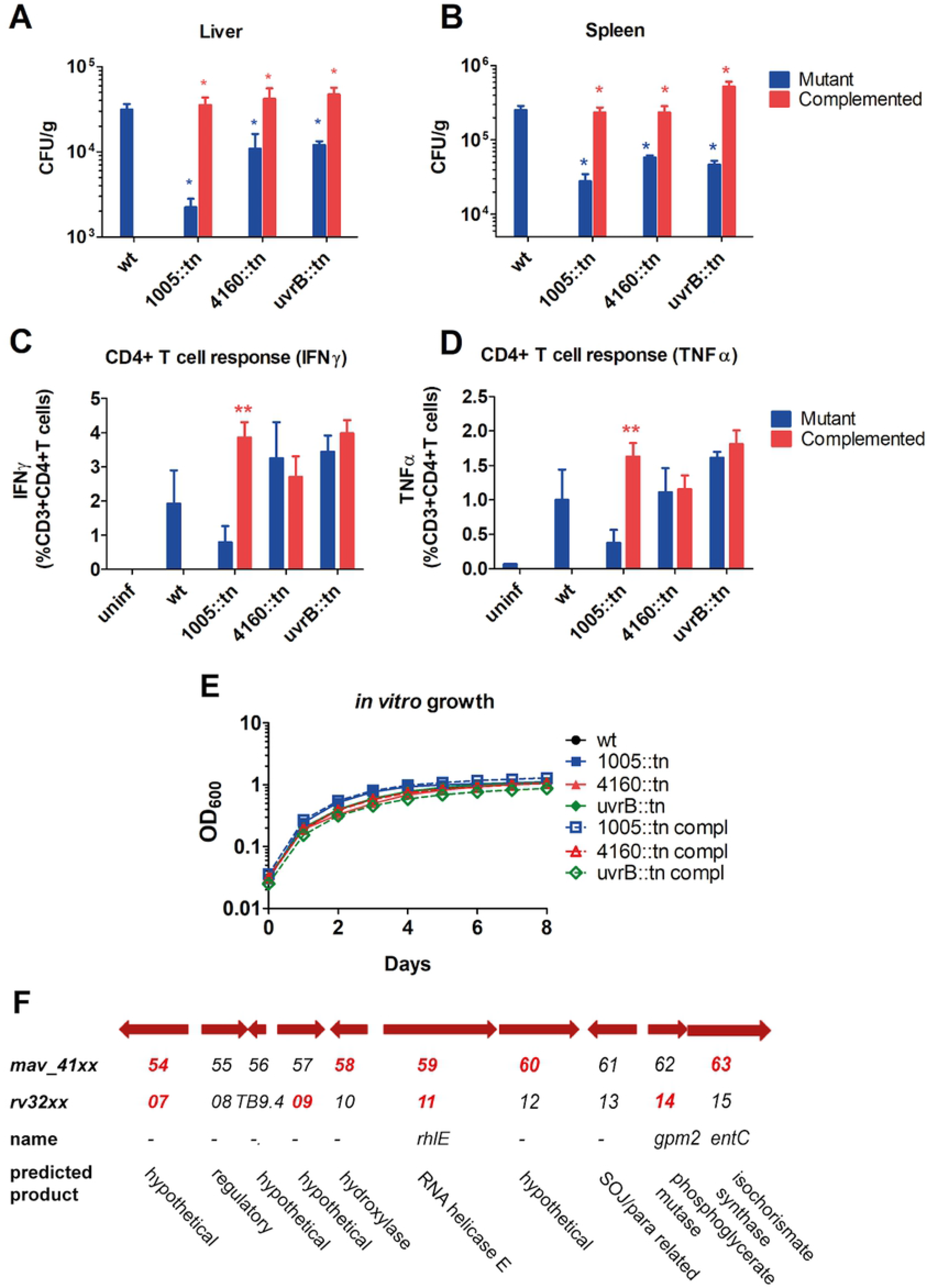
Validation of virulence genes. Mice were infected with MAH 11 wt or MAH 11 transposon insertion mutants with and without complementation. After 26 days of infection, the bacterial burden in spleen (A) and liver (B) was determined by CFUs per gram organ. Data show mean ± SEM of three or four infected mice in each group. *, P≤ 0.05; Mann Whitney U test, one-tailed, mutant compared to wt (blue asterisk) or mutant compared to complemented mutant (red asterisk). (C and D) MAH-specific CD4+ effector T cell response in mice infected with MAH 11 wt or MAH 11 transposon insertion mutants. 26 days post infection, splenocytes from all mice in each group were stimulated with MAH overnight and frequencies of IFNγ-producing (C) and TNFα-producing (D) CD4+ T helper cells were determined by flow-cytometry. Data show mean ± SEM. * P≤ 0.05, ** P≤ 0.01, and *** P≤ 0.001; unpaired students *t* test, two-tailed, compared to wt. (E) *in vitro* growth (7H9 medium) of MAH 11 wt and transposon insertion mutants with and without complementation. Data show mean ± SEM of three replicate samples per condition. The results are representative of three independent experiments. (F) Genetic region spanning from *mav_4154-4163*. Hits from our virulence screen (mav_41xx) and previously published *Mtb* virulence screen (Rv32xx) [16] are shown in bold red.

### A probable MFS transporter is required for MAH virulence

Next, we aimed to validate MAH determinants not previously implicated in mycobacterial virulence. A mutant of a probable MFS transporter, MAV_1005 (ortholog of Rv0876c), showed attenuated growth in our virulence screen. When we subjected an MAH 11 transposon mutant of this transporter (*1005::tn*) to mouse model infection, we saw a strong attenuation after 26 days of infection compared to wt and the complemented mutant (Fig 6A and B). Furthermore, the mutant but not the complemented mutant elicited a reduced MAH-specific CD4+ effector T cell response from the host (Fig 6C and D). The mutant did not show reduced fitness *in vitro* (Fig 6E). Taken together, our results suggest that *mav_1005* is crucial for MAH virulence.

### A hypothetical gene required for MAH virulence

Five genes located in close genomic vicinity (region spanning from *mav_4154* to *mav_4163,* Fig 6F) appeared as hits in our virulence screen. An insertion mutant of one of the genes, hypothetical gene *mav_4160* (*4160::tn*), was subjected to mouse model infection. After 26 days of infection, *4160::tn* showed attenuated growth in both liver and spleen (Fig 6A and B), though the MAH-specific effector CD4+ T cell response was not significantly reduced (Fig 6C and D). The mutant did not show reduced fitness *in vitro* (Fig 6E). Hence, our findings suggest that *mav_4160* is required for full MAH virulence.

### Inflammation and tissue pathology

We performed a broader characterization of the inflammation and tissue pathology in MAH 11-infected mice (including mutants and complemented mutants) at day 26 post infection. In brief, induction of organ homogenate cytokine production was low or not increased in response to infection (TNFα and IFNγ), except for IL-1β, which largely reflected organ bacterial loads (Fig 6 and S4 Fig). The overall low induction levels were not surprising; cytokines could be secreted by subsets of immune cells and act in an autocrine/paracrine manner e.g. in tissue granulomas. We have previously characterized the C57Bl/6 infection of MAH 104 in great detail [12]. Similar to what we observed with MAH 104, MAH 11 infection induced organ pathology seen as disruption of splenic pulp structures, infiltration of immune cells, inflammatory foci (granulomas) and giant cell formation (S5 Fig). However, no obvious differences in cytokine production or tissue pathology were seen between mutants and wt, nor mutants and complemented mutants. Nevertheless, the overall organ pathology and IL-1β-production grossly reflected bacterial loads.

## Discussion

To study the role of genes by loss of function is a powerful approach to understand how pathogens proliferate and avoid host elimination. We identified a virulent clinical isolate of MAH susceptible to genome-wide high-density gene disruption by ϕMycoMarT7-mediated transposon mutagenesis. The generated transposon library enabled us to define the MAH *in vitro* essential and virulence gene sets using a top-down discovery-based deep sequencing approach. A total of 674 genes were identified as required for normal growth *in vitro* (15% of total genes, similar to the proportion of required genes in *Mtb* [55]). There was a substantial overlap (71%) of *in vitro* essentials between MAH and *Mtb*, as well as many common virulence genes (e.g. *uvrABC*, *secA2*, *icl*) required for survival in a mouse model of infection. However, the majority of the virulence genes we identified were novel relative to *Mtb* TnSeq virulence screening [16]. Surprisingly, even with 1052 genes with no clear *Mtb* ortholog, only three of the MAH virulence genes were specific to MAH (i.e. did not have a mutual ortholog in the H37Rv genome). Two of these were PPE family proteins (*mav_4273/4*, found in MAC species but not *Mtb* [62]), which are unique to mycobacteria, associated with virulence, and some have been shown to be secreted by the ESX-5 secretion system in *M. marinum* and *Mtb* [65, 66]. In fact, we observed six genes within the MAH *esx-5* gene cluster among our virulence genes (*mav_2916-2933*), strongly indicating a crucial role of ESX-5 during MAH infection. However, when we subjected an ESX-5 mutant (*eccA5::tn*) to mouse infection, we did not see attenuated growth compared to wt (S6 A, B and E Fig), perhaps due to insufficient disruption of gene function in this mutant (transposon insertion in C-terminus), or that this particular gene is not required for ESX-5-mediated virulence in mice, as previously seen for *Mtb* [67]. Intriguingly though, we saw an increased CD4+ T cell host response when *eccA5* was overexpressed (S6 C and D Fig). Even so, it is possible that the two MAC-specific PPE proteins are secreted via the ESX-5 secretion system of MAH. These PPE proteins could be excellent candidates for targeted drug, vaccine and/or diagnostics discovery for improved control of MAC infections.

MAH and *Mtb* are both able to persist in human macrophages; however, *Mav* is environmental and opportunistic, while *Mtb* is an obligate human pathogen. The relatively modest overlap between MAH and *Mtb* in mutually orthologous virulence genes (46%) compared to the overlap of *in vitro* essential genes (71%) might reflect different mechanisms of virulence. However, interestingly, many of the genes that were unique to our MAH virulence screen have been experimentally proven required for *Mtb* virulence (exemplified by proteasome subunits *prcA*, *prcB* and genes encoding the *esx-5* secretion system [66, 68]). This suggests that a portion of the genes we found unique to MAH virulence, and that have *Mtb* mutual orthologs, might be required for *Mtb* virulence as well. Even so, it is evident that when MAH and *Mtb* are subjected to the same selective conditions (*in vitro* growth on 7H10 agar or *in vivo* growth in C57BL/6 mice), MAH depends on very few genes that do not have mutual orthologs in *Mtb* (~4% and ~3%, respectively). Whether this is true also for other MAH isolates, remains to be investigated. None of the genes within the major MAH 11 insertions (relative to MAH 104, listed in Figure 3B) were required for infection, and only four were essential *in vitro*. It has been shown that *M. marinum* customizes its virulence mechanisms to infect different animal cells [19]. It is thus possible that, if subjected to infection of other animal models, a greater proportion of genes specific to MAH would be required.

Interestingly, we identified genes on the two plasmids, pMD1 and pMD2, that were required for growth *in vitro* or *in vivo*. The *in vitro* growth defect seen in disruption of plasmid replication (*rep*) and partition (*parB*) genes might indicate that the presence of the two plasmids is required for efficient MAH proliferation, or that disturbing plasmid replication/partition reduces the global fitness of the MAH cells. It is currently unclear which role the three plasmid genes we found required *in vivo* might play during infection.

Using defined transposon mutants from our organized (plated) library, we validated a subset of our MAH virulence hits. We verified that the excision repair protein UvrB, a probable MFS transporter, and a hypothetical gene located within a genomic region of several identified virulence genes, are required for full MAH virulence. UvrB has previously been implicated in mycobacterial virulence [57], while the MFS transporter (*mav_1005*) and the hypothetical gene (*mav_4160*) were first validated as mycobacterial virulence factors by us. Of the six other virulence genes identified (based on MAH and *Mtb* screening) in the immediate vicinity of *mav_4160*, two encode hypothetical proteins, one a hydroxylase, one RNA helicase E (RhlE), one isochorismate synthase (EntC) and one phosphoglycerate mutase (Gmp2) (Fig. 6E). EntC was shown to be required for siderophore production in *M. smegmatis* [69], and might thus play a role in iron acquisition during infection. However, the mechanism of the MFS transporter, as well as *mav_4160* and the surrounding genes, in mycobacterial virulence remain to be elucidated.

*Mav* isolates exhibit high genetic variation [14, 42, 70, 71]. In accordance, we registered several large-scale insertions and deletions when we compared the genomic sequence of MAH 11 to MAH 104. The movement of genetic material between organisms is mediated by, among other mechanisms, phage transduction, natural transformation, and plasmid conjugation [72]. The cause (and effect) of *Mav* genomic plasticity is largely unknown. However, recently, mycobacterial conjugative plasmids have been identified, also in *Mav* [49, 50]. Interestingly, one of MAH 11’s plasmids, pMD2, is almost identical to previously described *Mav* plasmid pMA100 [49]. pMA100 was shown to transfer via conjugation between the slow growing mycobacteria *Mav* and *M. kansasii* in a mixed infection patient [49]. pMD2 might thus partake in genetic exchange between MAH 11 and other mycobacteria.

In conclusion, we identified a highly transformable MAH strain susceptible to ϕMycoMarT7-mediated transposon mutagenesis. This strain enabled genome-wide identification of *in vitro* essential and virulence genes in this species. Based on our screens, we identified growth and virulence genes specific to MAH, as well as shared with *Mtb*. MAH-specific genes might be excellent targets for MAC disease control, while shared genes might target related mycobacterial diseases as well. We validated two novel MAH genes required for infection. Since MAH 11 is used as a screening strain in mycobacterial drug discovery programs [36, 37], a comprehensive understanding of the genetic requirement for growth and infection of this strain is a direct asset for current initiatives towards new anti-mycobacterial therapies.

## Materials and methods

### Strains and growth conditions

MAH strains used in this study were MAH 104 and MAH 11 (NCBI GenBank accession number CP000479 and NBAW00000000, respectively). MAH strains were cultured in Middlebrook 7H9 (BD Difco) supplemented with 0.2% glycerol, 0.05% Tween 80 and 10% ADC (50 g BSA fraction V, 20 g dextrose, 8.5 g NaCl, 0.03 g catalase, dH_2_O up to 1 L) for liquid growth and in Middlebrook 7H10 (BD Difco) supplemented with 0.5% glycerol and 10% ADC for solid growth. For selection of transposon mutants, 20 μg/ml kanamycin and 0.1% Tween 80 was added to the 7H10 agar plate, the latter to simplify library harvest.

### Genome sequencing and annotation

DNA from late log phase cultures of MAH 11 was extracted using a Masterpure DNA Purification kit (Epicentre), prepared using the TruSeq genome DNA sample preparation kit (llumina, Inc.), and sequenced on an Illumina HiSeq 2500 instrument in paired-end mode with a read length of 125 bp. The genome sequence was assembled using a comparative assembly method, using MAH 104 as a reference sequence. SNPs were identified from mapped reads, which were aligned to the reference sequence using BWA [73] allowing up to 5/125 mismatches, and insertions/deletions (indels) were detected and repaired using local contig-building (as described in [74]). In addition, large-scale insertions and plasmid sequences were assembled de novo using Newbler (Roche, Inc.), and integrated into the genome where connectivity was supported by evidence from paired reads. The genome was annotated using PGAP [48], the NCBI Prokaryotic Genome Annotation Pipeline, which employs GeneMarkS+ to identify ORFs (along with RNAs and pseudo-genes) and assigns putative functions and gene identification based on homology. The sequences have been deposited in NCBI Genbank with accession numbers NBAW00000000 (aviumMD30 assembly of MAH 11), CM009838.1 (pMD1) and CM009839.1 (pMD2). aviumMD30 is the assembly and annotation analyzed in this paper. For genome completion, the MAH 11 strain was re-sequenced on a PacBio Sequal instrument. A total of 650Mb of long reads (up to 40kb) was collected. There were 130,570 subreads, with a mean read length of 4,937. Reads were mapped to the aviumMD30 assembly using blasr (version 5.3.2) [75]. Coverage of aligned segments was tabulated at each site. The PacBio data was used to confirm the connectivity of the genome by showing that all sites with low coverage (0-10x) in the Illumina data were spanned by PacBio reads, verifying that the chromosome consists of a single 5.1Mb contig. Similarly, the PacBio reads were aligned to the plasmid sequences pMD1 and pMD2 to confirm their continuity as circular, extra-chromosomal DNA. The updated assembly of the MAH 11 genome sequence (aviumMD36) is deposited in Genbank under accession number CP035744.

### Transformation

Competent MAH cells were prepared as previously optimized for *Mav* [33]. 100 μl of competent cells was electroporated with 2 μg plasmid DNA (pMSP12∷cfp, kind gift from Christine Cosma and Lalita Ramakrishnan [76]) in a 2 mm cuvette at settings 2.5 kV, 1000 ohm, and 25 μF. Cells were recovered overnight and plated at serial dilutions for colony forming unit (CFU) counts. CFU counts of pMSP12∷cfp-transformed bacteria selected on 7H10 with 20 μg/ml kanamycin were normalized to the CFU counts of transformed bacteria titered on 7H10 without antibiotics.

### Growth curves

MAH wt and mutants were grown in 7H9 medium until they reached stationary phase, then diluted to OD_600_ 0.02 in triplicates of 200 μl 7H9 medium in microplate honeycomb wells (Oy Growth Curves Ab Ltd.). The growth was monitored over the course of 10 days in a Bioscreen growth curve reader (Oy Growth Curves Ab Ltd.), shaking at 37 °C.

### Generation of MAH transposon mutant library

The MAH high-density transposon mutant library was prepared using ϕMycoMarT7 as previously described for *Mtb* [77], with the exceptions of growing the bacterial culture to stationary as opposed to exponential phase prior to transduction. The amount of ϕMycoMarT7 added for transduction was increased coordinately with the increased bacterial density. Both the phagestock and bacterial culture were heated to 37°C before transduction. The library was incubation at 37°C on 7H10 plates with Tween 80 (0.1%) and kanamycin (20 μg/ml) for 2-3 weeks.

### Transposon insertion sequencing (TnSeq)

The transposon library was harvested and pooled by scraping 7H10 plates with 170 000 colonies. Total DNA was purified using Masterpure DNA Purification kit (Epicentre), and prepared for TnSeq by PCR amplification of transposon:genome junctions and adapter ligation following the protocol in [53]. The samples were sequenced on an Illumina GAII instrument, collecting around 10 million 54 bp paired-end reads per sample. The reads were processed using TPP in Transit [54], which counts reads mapping to each TA dinucleotide site (after eliminating reads sharing the same template barcode; [53]).

### MAH *in vitro* essential gene set

Essential genes were identified using a Hidden Markov Model (HMM, [78]), incorporated into Transit [54]. The HMM is a Bayesian statistical model that parses the genome into contiguous regions labeled as one of 4 states - essential (ES), non-essential (NE), growth-defect (GD, suppressed insertion counts), or growth-advantaged (GA, insertion counts higher than average) - based on local insertion density and mean read count at TA sites. The output of the HMM was processed by labeling each gene with the majority state among the TA sites spanned by it.

### Mouse infection

For MAH 104 and MAH 11 infection experiments, groups of four C57BL/6 mice were infected *intraperitoneally* with 5×10^7^ CFU/mouse as previously described [12]. On day 22 and day 50 post infection, MAH-specific effector T cell responses and bacterial load were analyzed. Numbers of CFUs per gram organ were measured by plating serial dilutions of spleen and liver homogenates on 7H10 plates. For the virulence gene screen, six mice were infected *intraperitoneally* with 6×10^7^ CFU/mouse of the MAH 11 transposon mutant library. After 22 days of infection, mice were sacrificed and the liver and spleen were harvested, homogenized and plated on 7H10 (livers and spleens from two mice were pooled to make one library, resulting in three liver and three spleen libraries in total from the six mice). After 2-3 weeks at 37°C, the colonies were scraped and DNA prepared for sequencing as described above for TnSeq. For *in vivo* validation experiments of virulence genes, groups of 4-5 C57BL/6 mice were infected *intraperitoneally* with (approximately) 7.5×10^7^ CFU/mouse MAH 11 wt or MAH 11 transposon insertion mutants. On day 26 post infection, bacterial load in liver and spleen, MAH-specific effector T cell responses, cytokine levels in organs and serum and histopathology were analyzed.

### MAH-specific T cell response

Isolated splenocytes from infected mice were stimulated overnight with MAH (MOI 3:1); protein transport inhibitor cocktail (eBioscience) was added for the last 4h of incubation. Unstimulated cells were used as controls. Cells were harvested and stained with Fixable Viability Dye eFluor 780 (eBioscience) and fluorescence-labelled monoclonal antibodies against CD3 (FITC, eBioscience) and CD4 (Alexa Fluor700 or Brilliant Violet 605, both from BioLegend). After fixation and permeabilization, intracellular cytokine staining was performed with fluorescent monoclonal antibody against IFNγ (Phycoerythrin, eBioscience) and TNFα (Allophycocyanin, BioLegend). Cells were analyzed by flow cytometry on a BD LSR II flow-cytometer (BD Biosciences) and data subsequently analyzed with FlowJo (FlowJo, LLC) and GraphPad Prism (GraphPad Software, Inc.) software. Frequencies of IFNγ- and TNFα-producing CD4+ effector T cells were analyzed from FSC/SSC-gated, viable CD3+CD4+ T cells. The method is described in further detail in [12].

### Cytokine measurements

IL-1β, IFN-γ and TNF-α levels were analyzed in serum as well as spleen and liver homogenates from infected mice using a custom-made ProcartaPlex immunoassay panel (Affimetrix, eBioscience) according to the manufacturer’s protocol.

### Histopathology

Standard hematoxylin & eosin staining of spleen and liver sections was performed at the Cellular and Molecular Imaging Core Facility (CMIC) at NTNU as described previously [12]. Images were acquired with a Nikon E400 microscope and NIS-Elements BR imaging software (Nikon Instruments, Melville, NY, USA).

### MAH virulence gene set

For comparative analysis between the *in vitro*-selected and the *in vivo* (mouse infection)-selected transposon libraries, the TRANSIT-incorporated ‘resampling’ algorithm was used [54]. Resampling is analogous to a permutation test, examining if the sum of transposon insertion read counts differs significantly between conditions.

### Bulk-identification of transposon insertion sites in the organized MAH library

#### Organizing, pooling and sequence-tagging the library

9216 colonies were picked and transferred to 24 384-well plates in duplicate using a Genetix QPixII colony picker and software QSoft XP Picking. The cultures were incubated at 37°C for 3 weeks. The library was pooled by plates (24), columns (24) and rows (16), giving a total of 64 culture pools using a Freedom EVO 200 (TECAN) liquid handling robot and software EVOsim. Of the total 64 pools, each well should be represented three times; in one plate pool, in one column pool and in one row pool. The various pools were tagged by ligating barcoded adapters to the DNA fragments after DNA purification. Barcodes 1-24 represent plates and columns 1-24, while barcodes 1-16 represent rows 1-16 (further details in S1 Materials and Methods and S3 Table).

#### Analysis of arrayed library of Tn mutants

The three tagged pools were sequenced on an Illumina HiSeq 2500 with 125 bp paired-end reads, collecting 3.2-5.3 million pairs of reads each pool. The genomic portions of the reads (in read 1) were mapped to TA sites in the MAH 11 genome (including plasmids) using BWA [73]. The barcodes (8 bp embedded in read 2) were extracted and tabulated for each insertion coordinate. Subsequently, a script was written that compared the TA sites represented by each combination of row-, column-, and plate-barcode to associate each TA site with the most probable well. The count of each barcode for each site was normalized by dividing by the barcode total over all sites. Barcodes with insufficient counts (<10,000 total) were excluded from the calculation. Then the relative frequencies of a plate-, row-, and column-barcodes for each site, 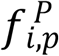, 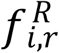 and 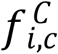 were computed by dividing by the total abundance of plate-, row-, and column barcodes represented by the site. Finally, a score 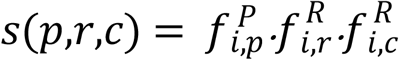 was computed for each possible combination of plate, row-, and column-barcodes that represented the likelihood of the well assignment for each TA site *i*. Wells with high probability (*s*≥0.8 for the maximal combination) were accepted as unique assignments; wells with *s*<0.8 were rejected as ambiguous (i.e. potentially mapping to multiple wells).

### Verification of mutants by PCR and Sanger sequencing

Total DNA was isolated using Masterpure Complete DNA Isolation kit (Epicentre). PCR to confirm presence of transposon was preformed using GoTaq^®^ Green Master Mix (Promega) and primers KanF (TGGATTGCACGCAGGTTCTC) and KanR (CGTCAAGAAGGCGATAGAAG). Sanger sequencing was preformed using gDNA as template prepared with BigDye Terminator Cycle Sequencing kit v.1.1 (Thermo Fisher); with 60 rounds cycles of 95°C 30 sec, 52°C 30 sec, 60°C 4 min, and primer KanSeq2 (CTTCCTCGTGCTTTACGG) reading directly into the gDNA. Samples were purified using BigDye XTerminator kit (Applied Biosystems) and sequenced on an ABI130xl.

### Complementation of transposon insertion mutants

Plasmids for complementation of transposon insertion mutations were constructed by cloning the wt version of the disrupted gene into the mycobacterium-escherichia coli shuttle vector pMV261 ([79]) for constitutive expression. To select for the plasmids upon transformation into the mutated strains, the kanamycin resistance gene from pMV261 was swapped with the hygromycin resistance gene of pUV15TetORm [80], creating pMV261H. *b6k05_04950* and *b6k05_04945*, *b6k05_18820*, *b6k05_12440*, and *b6k05_13510* were amplified from the MAH 11 genome and cloned into pMV261H, resulting in plasmids pMV261H *1005*, pMV261H *4160*, pMV261 *eccA5*, and pMV261 *uvrB*, respectively. All clonings were performed using Gibson Assembly^®^ Master Mix (New England Biolabs). Primer sequences can be provided upon request. The complementing plasmids were transformed into their respective MAH 11 mutant, resulting in strains *1005::tn* compl., *4160::tn* compl., *eccA5::tn* compl., and *uvrB::tn* compl.

### Ethics statement

The protocols on animal work were approved by the Norwegian Animal Research Authorities (Forsøksdyrutvalget, FOTS ID 5955). All procedures involving mice experiments were carried out in accordance with institutional guidelines, national legislation and the Directive of the European Convention for the protection of vertebrate animals used for scientific purposes (2010/63/EU).

## Acknowledgements

We thank Jun-Rong Wei, Jung-Yien Chien and Po-Ren Hsueh for providing clinical MAH isolates from the National Taiwan University Hospital, the Medical Genetics Department at St. Olavs Hospital for assistance with Sanger sequencing, the Cellular and Molecular Imaging Core Facility (CMIC) at NTNU for histopathology staining and personnel at the Comparative Medicine Core Facility (CoMed) at NTNU for assistance in animal experiments. CMIC and CoMed are funded by the Faculty of Medicine and Health Science at NTNU and Central Norway Regional Health Authority. We acknowledge the help of the Texas A&M Sequencing Center (Genomics & Bioinformatics Service, Dr. Charlie Johnson, director).

## Supporting information

**S1 Fig. Verification of transposon insertions.** 10 colonies were picked from a plated MAH library and subjected to PCR using primers specific for the *Himar1* transposon. Positive control (pos ctrl) was ϕMycoMarT7, negative control (neg ctrl) was un-transduced MAH 11. Colony number 4 gave a band in the expected size when resubjected to PCR.

**S2 Fig. MAH phylogenetic tree.** A phylogenetic tree containing MAH 11, MAH 104 and 19 other *M. avium* strains was constructed using PHYLIP. The subspecies MAH, MAA, and MAP, were confirmed by their hsp60 sequence.

**S3 Fig. MAH cell wall integrity assay.** Sensitivity to SDS was assessed by spotting dilutions of MAH 104 and MAH 11 (and transposon mutants) on 7H10 plates prepared with or without 0.01% SDS.

**S4 Fig. Cytokine production during MAH 11 mouse infection.** Mice were infected for 26 days with MAH 11 wt or MAH 11 transposon insertion mutants with and without complementation. Levels of IL-1β (A), TNFα (B) and IFNγ (C) were analyzed in spleen and liver homogenates and serum. Data show mean ± SEM of three or four infected mice in each group. The dotted line represents the cytokine level of uninfected mice.

**S5 Fig. Histopathology during MAH 11 mouse infection.** Mice were infected with MAH 11 wt or MAH 11 transposon insertion mutants with and without complementation. After 26 days, hematoxylin & eosin staining was performed on spleen and liver sections. The panel shows representative 10x and 40x magnification images of spleen and liver from one out of three or four infected mice in each group.

**S6 Fig. Mouse infection with MAH 11 eccA5::tn ESX-5 mutant.** Mice were infected with MAH 11 wt or *eccA5::tn* mutant with and without complementation (4 mice in each group). After 26 days of infection, the bacterial burden in spleen (A) and liver (B) was determined by CFUs per gram organ. (C and D) MAH-specific CD4+ T cell response in mice infected for 26 days with MAH 11 wt or *eccA5::tn* mutant with and without complementation. Splenocytes from all mice in each group were stimulated with MAH overnight and frequencies of IFNγ- (C) and TNFα- (D) producing CD4+ T helper cells were determined by flow-cytometry. Data show mean ± SEM. * P≤ 0.05; unpaired students *t* test, two-tailed, compared to wt. (E) *in vitro* growth (7H9 medium) of MAH 11 wt and eccA5::tn mutant with and without complementation. Data show mean ± SEM of three replicate samples per condition. The results are representative of three independent experiments.

**S1 Table. Output statistics for the sequenced MAH libraries.**

**S2 Table. Verification of transposon insertion sites by Sanger sequencing.**

**S3 Table. Adapters used in TnSeq of the organized MAH library.**

**S1 Materials and Methods. Bulk-identification of transposon insertion sites in an organized MAH library.**

### S1 Datasets (A-O)

A – MAH 104 and MAH 11 mutual orthologs.
B – MAH 11 *in vivo* genetic requirement and MAH 11 and *Mtb* mutual orthologs.
C – MAH 11 and *Mtb* best orthologs.
D – Plasmid pMD1 annotation.
E – Plasmid pMD2 annotation.
F – pMA100 and pMD2 mutual orthologs.
G – MAH 11 *in vitro* genetic requirement.
H – pMD1 *in vitro* genetic requirement.
I – pMD2 *in vitro* genetic requirement.
J – pMD1 (spleen) *in vivo* genetic requirement.
K – pMD1 (liver) *in vivo* genetic requirement.
L – pMD2 (spleen) *in vivo* genetic requirement.
M – pMD2 (liver) *in vivo* genetic requirement.
N – Location of transposons in MAH 11 organized library.
O – Map of mutants/well in MAH 11 organized library.

